# Behavioral Evidence for Two Modes of Attention

**DOI:** 10.1101/2024.09.12.612641

**Authors:** Akanksha Gupta, Tomas E. Matthews, Virginia B. Penhune, Benjamin Morillon

## Abstract

Attention modulates sensory gain to select and optimize the processing of behaviorally relevant events. It has been hypothesized that attention can operate in either a rhythmic or continuous mode, depending on the nature of sensory stimulation. Despite this conceptual framework, direct behavioral evidence has been scarce. Our study explores when attention operates in a rhythmic mode through a series of nine interrelated behavioral experiments with varying stream lengths, stimulus types, attended features, and tasks. The rhythmic mode optimally operates at approximately 1.5 Hz and is prevalent in perceptual tasks involving long (> 7 s) auditory streams. Our results are supported by a model of coupled oscillators, illustrating that variations in the system’s noise level can induce shifts between continuous and rhythmic modes. Finally, the rhythmic mode is absent in syllable categorization tasks. Overall, this study provides empirical evidence for two modes of attention and defines their conditions of operation.

## Introduction

In an ever-shifting environment with a constant influx of dynamic information, the human brain, through attentional sampling, orchestrates its limited resources to select and optimize the processing of behaviorally relevant events^1–3^. In human audition, speech and music present temporal regularities that aid in orienting the fluctuations of attention to the external rhythm^4–6^. This notion was initially put forward as the auditory dynamic attending theory^7,8^. One way dynamic attending could be implemented in the brain is the synchronization of neural activity to the temporal structure of attended events^9–11^, which temporally modulates the excitability of task-relevant neuronal populations, thereby enhancing perception^9,11–14^. Nonetheless, behavioral evidence that temporal regularities lead to improved processing of sensory events is mixed and unclear.

Several studies have demonstrated improved behavioral performance associated with auditory temporal regularity^15–20^. Within the context of beat-based timing tasks, both improved accuracy and faster reaction times have been shown when detecting stimuli present in a temporally regular stream compared to an irregular stream^18,19^. Furthermore, in similar experimental paradigms, perceptual performance is dependent on stimulus presentation rate, with optimal accuracy observed within the range of 1 to 2 Hz, peaking at approximately 1.5 Hz^20^. Notably, this is also the rate at which our strongest sense of musical beat occurs^21,22^. However, other studies have shown conflicting^23–26^ or null^27^ results regarding the positive behavioral impact of temporal regularities in similar beat-based auditory paradigms. These inconsistencies in experimental outcomes between research groups could notably arise due to differences in the stimulus (e.g., length of the sequence^26^) or the task (e.g., detection versus discrimination^23,26,27^). This crucially suggests that dynamic attending is not always in effect and may be irrelevant in some situations.

One proposal suggests that two fundamentally different modes of attention exist. Depending on the temporal structure of the external events, attention could operate in either a rhythmic or continuous processing mode^4^. The rhythmic attentional mode would engage when the stimulus has a behaviorally relevant temporal structure, typically periodic and predictable.

Conversely, the continuous mode would operate when the temporal structure is behaviorally irrelevant, random, or unpredictable^4,28^. However, empirical evidence of the existence of two modes of attention is scarce, and the specifics of when these two modes operate have yet to be explored^4,29,30^.

Here, we investigated which key variables could lead to these two attentional modes, and we hypothesize that differences in the stimulus or task structure explain the conflicting results in the literature concerning rhythmic facilitation of auditory perception. To this end, we designed 9 behavioral paradigms in which we varied the stream length (short or long), the presence of distractors (yes or no), the type of stimulus (pure tones or syllables), the attended feature (pitch or duration), and the task (perceptual or categorization). For each experiment, we investigated the impact of stimulus presentation rate on behavioral performance, the rhythmic mode being characterized by an optimal perceptual enhancement at approximately 1.5 Hz, at least for perceptual tasks involving tones or music^20,22^. Finally, in contrast to tones or music perception, speech perception is optimal at syllable rates of 2 to 8 Hz^31^, with a peak at approximately 4.5 Hz^5,31–335,31^. Nevertheless, why different optimal rates are reported for speech and music processing, and what drives this difference is unclear. By investigating different stimulus types (e.g., syllables) and tasks (e.g., categorization), we aimed to further generalize these results, thereby contributing to a broad understanding of the two modes of attention.

Our findings reveal a rhythmic mode of operation —characterized by an optimal processing rate of approximately 1.5 Hz— during perceptual tasks demanding more attentional resources, namely the detection of deviant targets in long auditory streams. Remarkably, this mode of operation remains consistent across different stimuli (pure tones or syllables) and different attended features (pitch or duration). Our results can be explained by a model of coupled oscillators, where noise in the system gradually decreases over time, facilitating the transition to a rhythmic mode of operation, after approximately 7 s. Finally, this rhythmic attentional mode does not manifest during the categorization of syllables, highlighting its specificity to perceptual tasks.

## Results

In this study, we tested the influence of stream length, presence of distractors, attended feature, stimulus type, and task type on the emergence of a continuous or rhythmic attentional mode of operation. Participants engaged in two paradigms: an ongoing detection task (with streams lasting between 30 and 70 s, mean = 49 s) and a Yes/No task (with streams lasting between 3 and 12 s, mean = 7 s). In ongoing detection tasks, participants pressed the key whenever a deviant target was presented (each stream contained a total of 8 or 9 deviants), whereas, in Yes/No tasks, they decided at the end of the stream if a single deviant target was present or not in the stream. The difficulty level of each experiment was individually titrated using a psychophysical staircase at a tempo of 3 Hz (except for 1.5 Hz in experiment 3). In all experiments, the tempi varied from 0.5 to 6.5 Hz (except for 0.7 to 3.5 Hz in experiment 3).

### The length of the stream determines the attentional mode

In Experiments 1 and 2, we employed pure tone stimuli to investigate pitch discrimination performance using isochronous tempi ranging from 0.5 - 6.5 Hz, with a long stream ongoing detection and a short stream Yes/No tasks (Fig. 1a and 1c; see Methods). The average difficulty level (pitch deviation rate) was 2.45% (Mdn = 2.35, SD = 1.16) for Experiment 1 (Supplementary Fig. 1a) and 7.3% (Mdn = 7.92, SD = 2.02) for Experiment 2 (Supplementary Fig. 1c).

**Fig. 1:**
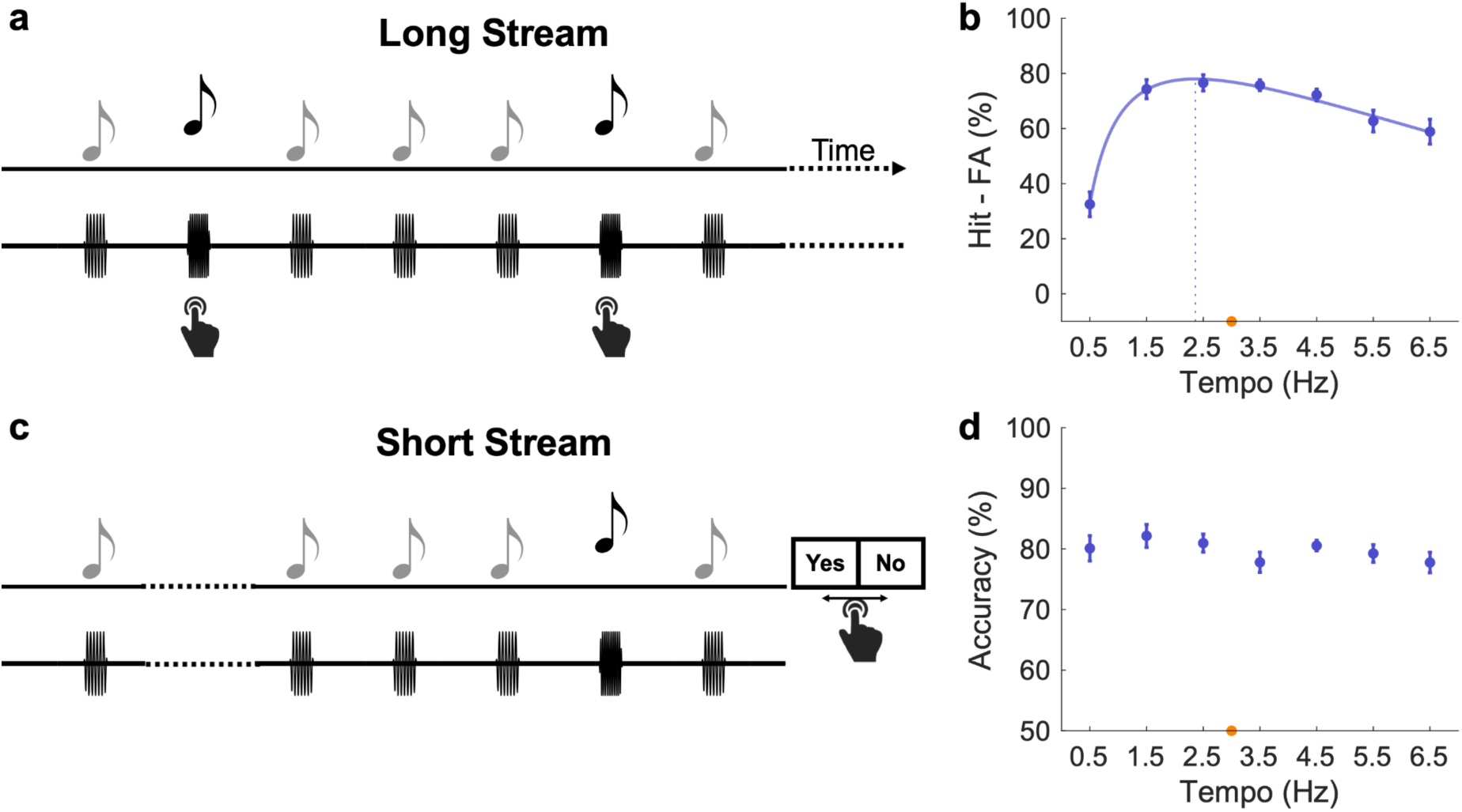
Experiments 1-2. Detection of higher-pitched tones. **a** Each trial consisted of a long periodic stream (∼1 min) of pure tones. The rate of presentation of the stimuli (tempi) varied across conditions (0.5 Hz to 6.5 Hz, in 1 Hz steps). Participants performed an ongoing detection task of perceptual deviants corresponding to higher pitch tones. **b** Average performance (Hit minus False alarm (FA) rate; blue dots) per condition (*n* = 20). **c** Each trial consisted of a short periodic stream (<10 s) of pure tones. The tempi varied across conditions (as in **a**). Participants performed a Yes/No task and indicated the presence or absence of a perceptual deviant (higher-pitched tone) at the end of each trial. **d** Average performance (Accuracy; blue dots) per condition (*n* = 21). **(b, d)** If performance significantly differed between conditions (repeated-measures ANOVA), data were approximated with a polynomial function (plain line) and an optimal frequency (leading to a maximal performance) could be estimated (vertical dashed line). The orange dot indicates the tempo (3 Hz) that served to assess the individual difficulty level. Error bars indicate ±SEM.

In Experiment 1, the comparison of performance (Hit minus False alarm (FA) rate) across different tempi revealed significant differences (repeated-measures ANOVA: F(6,114) = 17.47, p < 0.001; Fig. 1b). The performance profile exhibited an inverse U-shaped pattern, approximated by a second-order log polynomial function (adjusted r2 = 0.99, see Methods). The local maximum of this function estimates the tempo at which performance is optimal, indicating an optimal frequency of 2.36 Hz.

Conversely, in Experiment 2, the comparison of accuracy rates across tempi did not reveal significant performance fluctuations (repeated-measures ANOVA: F(6,114) = 0.86, p = 0.50; Fig. 1d). In this case, there is no discernible optimal frequency.

This suggests that the length of the stream plays a crucial role in determining the operative attentional mode. We hence hypothesized that longer streams necessitate increased attention, thus leading to the activation of a rhythmic mode to improve perceptual accuracy, whereas the continuous mode is active for short streams, leading to similar response accuracy for all stream rhythms. Of note, we ruled out the possibility that the type of paradigm (ongoing detection or Yes/No tasks) determines these results (see below, ‘Control analysis’ section).

### A model of coupled oscillators to generalize the results

We explored the possibility of our results being explicable through a standard model of coupled oscillators^34^. To comprehend the influence of stream length leading to optimization, we constructed a basic model incorporating an auditory attention oscillator, characterized by an intrinsic rhythm susceptible to external synchronization. This results in a model featuring two interconnected phase oscillators (Stimulus (S) and Attention (A)) with adjustable time delays, noise, and coupling strength (Fig. 2a).

**Fig. 2:**
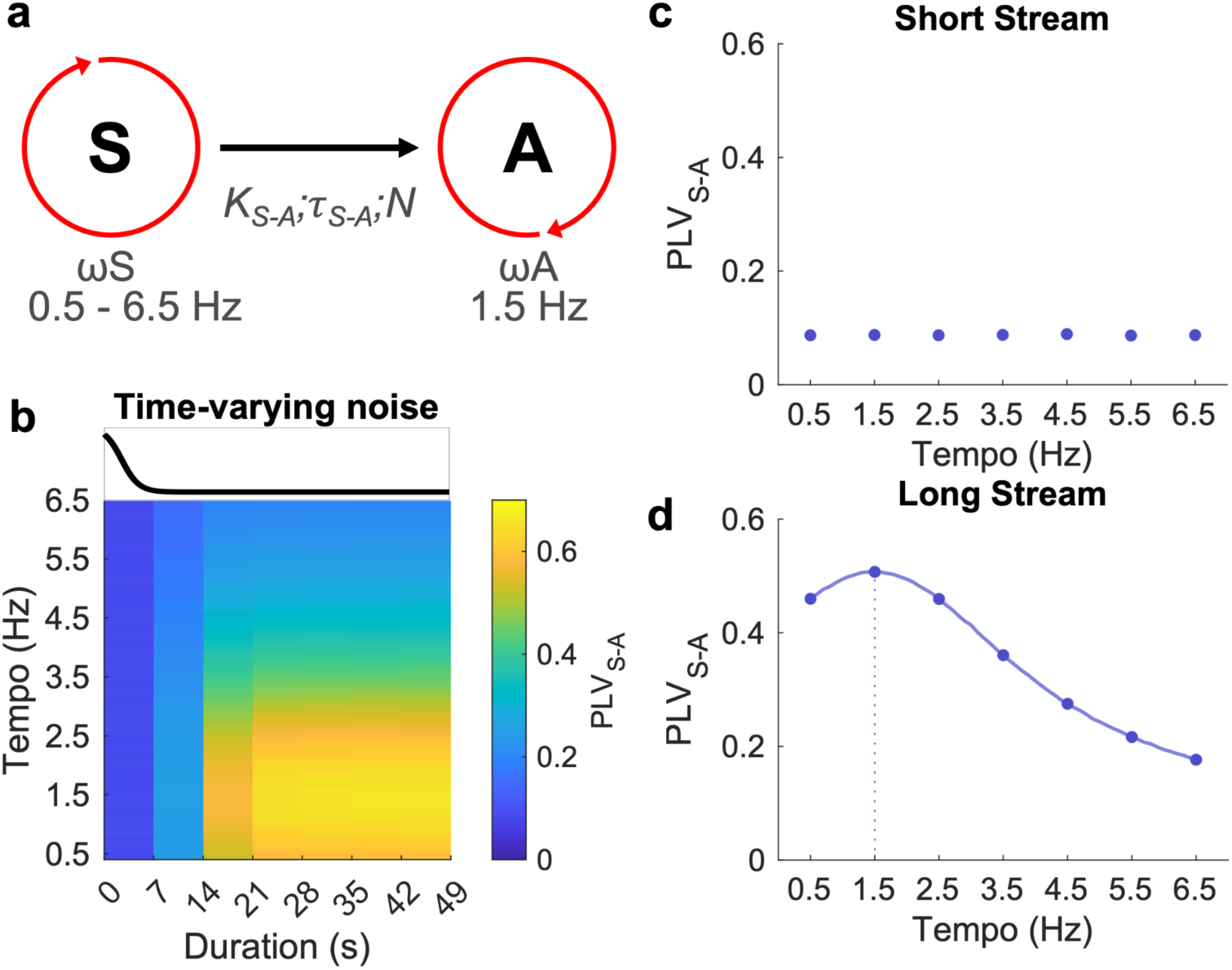
A model of coupled oscillators to generalize the results. **a** Model of two delay-coupled phase oscillators approximating the selective coupling between the external beat (stimulus S) and auditory-specific temporal attention (A). The external oscillator (S) influences the attention oscillator with a specific strength K, delay τ, and noise N. The phase-locking value (PLV) reflects the capacity of the attention oscillator (A) to synchronize with the external beat (S). It is thus used as an approximation of behavioral performance. **b** Heatmap depicting the evolution of PLV for each condition (tempi), from 0 to 49 s, in steps of 7 s. The tempi varied from 0.5 Hz to 6.5 Hz (in 0.01 Hz steps) and data of n = 20 participants were simulated. K and τ parameters were kept constant, and the noise N varied over time (see **b** inset). **(c, d)** Time-averaged PLV corresponding to **(c)** short (0-7 s) or **(d)** long (0-49 s) trials. If performance significantly differed between conditions (repeated-measures ANOVA), data were approximated with a polynomial function (plain line) and an optimal frequency (leading to a maximal performance) could be estimated (vertical dashed line).

The external stimulus (S) varied in frequency to emulate our various experimental conditions (ranging from 0.5 to 6.5 Hz). The internal frequency of the auditory attention oscillator (A) remained constant, representing the optimal sampling rate for temporal attention in the auditory domain at 1.5 Hz^20^. We then adjusted coupling strengths (K), time delays (τ), and the level of internal noise (D) individually with time, to align with the distinct behavioral outcomes. Behavioral performance was estimated using the phase-locking value (PLV) between the external beat (S) and the attention oscillator (A), serving as an indicator of the ability of the oscillator to synchronize with the external beat. We found that when the noise was adjusted using a reverse sigmoid function (Fig. 2b), such that the noise in the system gradually decreased over time, we could replicate the behavioral results (Fig. 2c, d). The comparison of performance (time-averaged PLV) across different tempi did not have significant performance fluctuations for short streams of 0 - 7 s (repeated-measures ANOVA: F(6,114) = 0.46, p = 0.78; Fig. 2c), while the performance for long streams of 0 - 49 s showed significant differences (repeated-measures ANOVA: F(6,114) = 44472.64, p < 0.001; Fig. 2d). Furthermore, the performance profile for long streams exhibited an inverse U-shaped pattern with an optimal frequency of 1.5 Hz. In contrast, varying the coupling strength (Supplementary Fig. 2a) and time delay (Supplementary Fig. 2b) linearly with time resulted in similar patterns of performance for both short and long streams (Supplementary Fig. 2c, d). This indicates that only variations of the noise within the model lead to the specific patterns we observed in the behavioral results.

### The presence of distractors necessitates optimization for short streams

In Experiment 3, we employed pure tone stimuli and added distractors in a short-stream pitch evidence-accumulation task (Fig. 3a; see Methods) using isochronous tempi ranging from 0.7 - 3.5 Hz. The average difficulty level (pitch deviation rate) was 42.81% (Mdn = 39.01, SD = 26.95) (Supplementary Fig. 3a). In this experiment, the task imposes a rhythmic mode of operation, i.e. above-chance performance can only be achieved if participants modulate their attention in time at the tempo, which necessitates some training^20,35,36^.

**Fig. 3:**
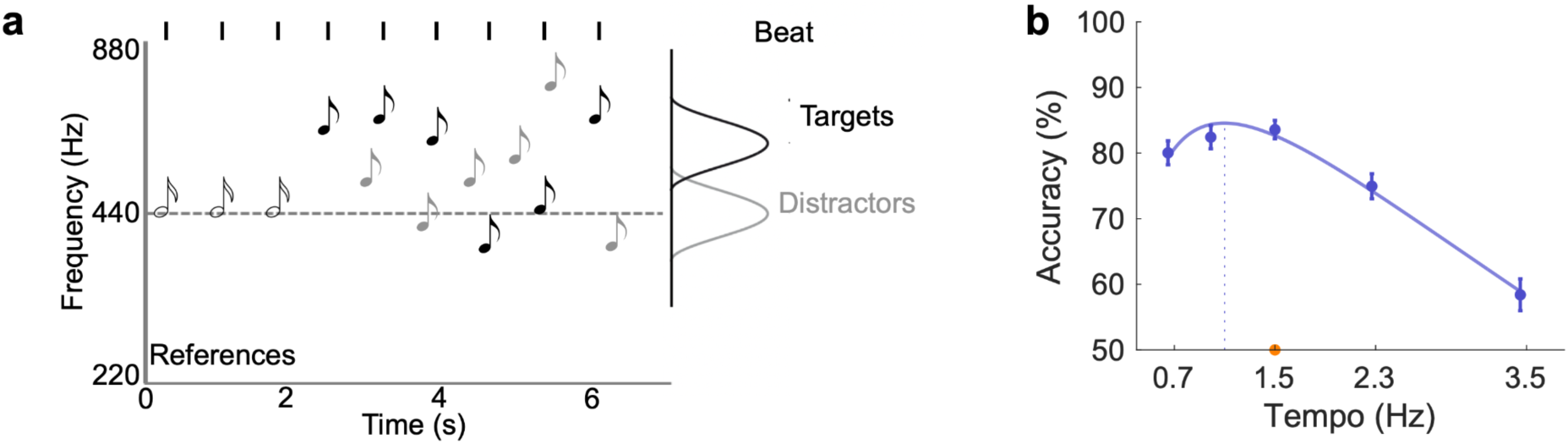
Experiment 3. Pitch evidence-accumulation. **a** Each trial consisted of a short periodic stream (<10 s) of 15 pure tones. Three reference tones preceded an alternation of six targets and six distractor tones of variable frequencies. Targets occurred in phase with the preceding references, whereas distractors occurred in antiphase. Participants had to decide whether the mean frequency of targets was higher or lower than the reference frequency. **b** Average performance (Accuracy; blue dots) per condition (*n* = 20). The tempi varied across conditions (0.7 Hz to 3.5 Hz). The same conventions as in Fig. 1 b-d.

In contrast to Experiment 2, which also had a short stream, we found a significant difference in performance accuracy across tempi (repeated-measures ANOVA: F(4,76) = 23.66, p < 0.001; Fig. 3b). Like in Experiment 1, the performance profile exhibited an inverse U-shaped pattern, approximated by a second-order log polynomial function (adjusted r2 = 0.97; see Methods), indicating an optimal frequency of 1.10 Hz. These results show that the rhythmic mode of operation can emerge rapidly (< 7 s) if the task imposes it.

### No influence of attended perceptual features on attentional modes

In Experiments 4 and 5, we employed pure tone stimuli to investigate —not pitch but— duration discrimination to replicate and generalize our results to different attended perceptual features (Fig. 4a and Fig 4c; see Methods). The average difficulty level (duration of deviant target compared to an 80 ms long standard) was 139.15 ms (Mdn = 138, SD = 22.06) for Experiment 4 (Supplementary Fig. 4a) and 108.2 ms (Mdn = 103, SD = 12.43) for Experiment 5 (Supplementary Fig. 4c).

**Fig. 4:**
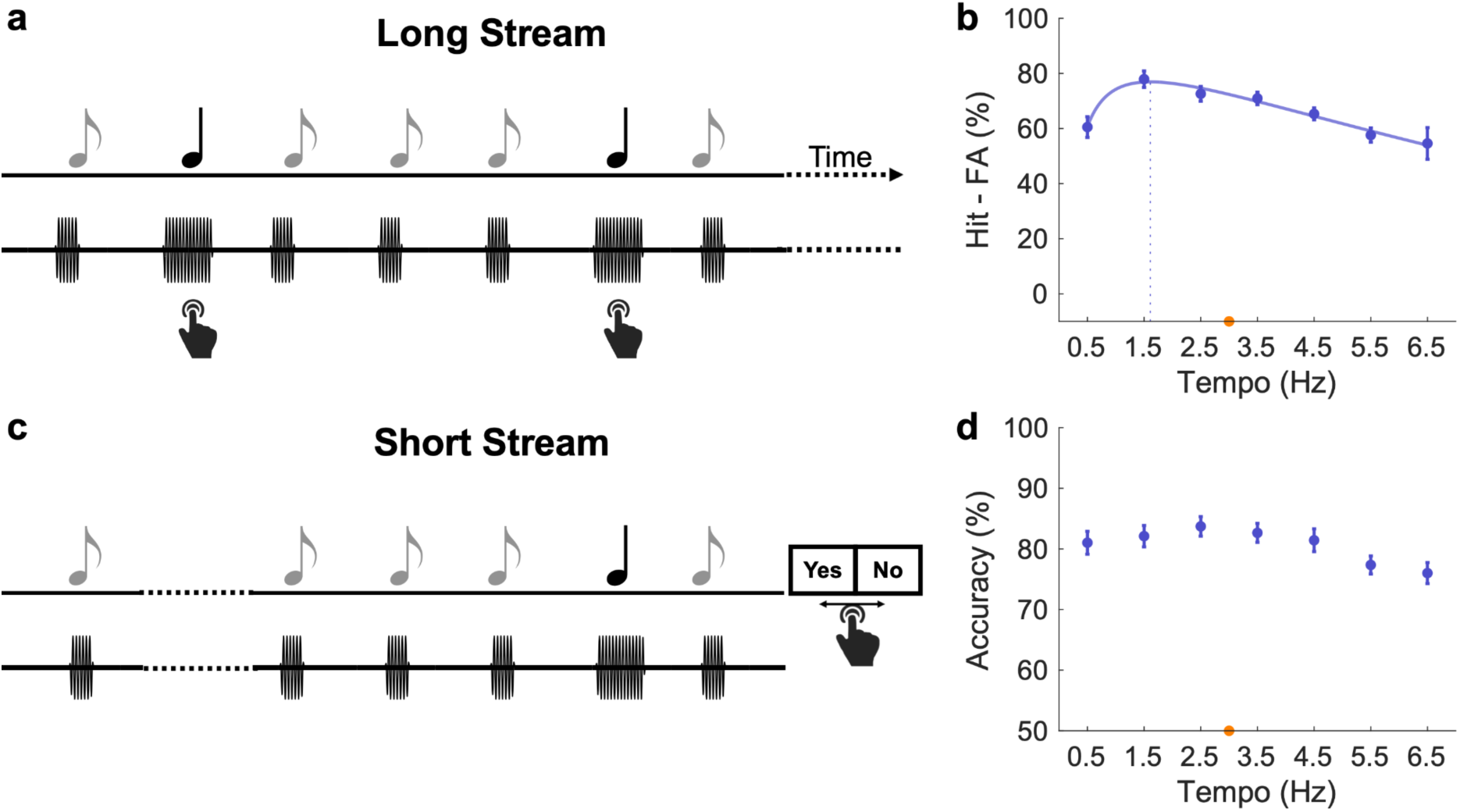
Experiments 4-5. Detection of longer tones. **a** Each trial consisted of a long periodic stream (∼1 min) of pure tones. Participants performed an ongoing detection task of perceptual deviants corresponding to tones of longer duration. **b** Average performance (Hit minus FA rate; blue dots) per condition (*n* = 20). **c** Each trial consisted of a short periodic stream (<10 s) of pure tones. Participants performed a Yes/No task and indicated the presence or absence of a perceptual deviant (longer tone) at the end of each trial. **d** Average performance (Accuracy; blue dots) per condition (*n* = 20). The same conventions as in Fig. 1 b-d.

In Experiment 4, the comparison of performance (Hit minus FA rate) across different tempi revealed significant differences (repeated-measures ANOVA: F(6,114) = 5.55, p = 0.006; Fig. 4b), with an inverse U-shaped pattern (second-order log polynomial function; adjusted r2 = 0.97) peaking at 1.61 Hz.

Conversely, in Experiment 5, the comparison of accuracy rates across tempi did not reveal significant performance fluctuations (repeated-measures ANOVA: F(6,114) = 2.38, p = 0.054; Fig. 4d). In this case, there is no discernible optimal frequency.

This indicates that the perceptual feature being attended to (pitch, duration) does not have an influence on which attentional mode is operational, only the stream length does.

### No influence of stimulus type on attentional modes

In Experiments 6 and 7, we asked participants to detect —not pure tones, but— higher-pitch syllables, to test the generalizability of our results to different stimulus types (Fig. 5a and Fig 5c; see Methods). The average difficulty level (pitch deviation rate; arbitrary units) was 6.84 (Mdn = 7, SD = 2.9) for Experiment 6 (Supplementary Fig. 5a) and 4.8 (Mdn = 4, SD = 3.39) for Experiment 7 (Supplementary Fig. 5c).

**Fig. 5:**
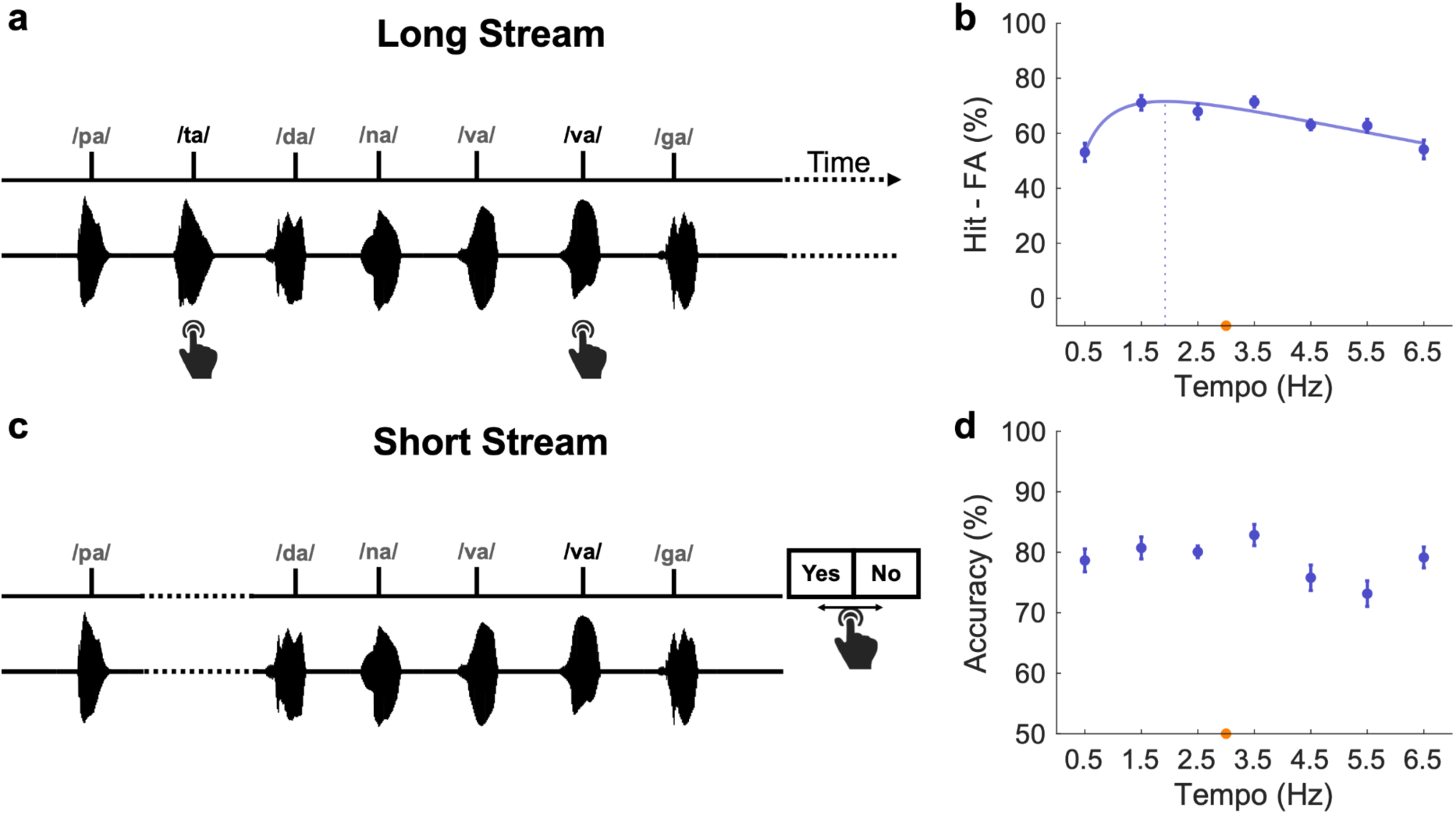
Experiments 6-7. Detection of higher-pitched syllables. **a** Each trial consisted of a long periodic stream (∼1 min) of syllables. Participants performed an ongoing detection task of perceptual deviants corresponding to syllables of higher pitch. **b** Average performance (Hit minus FA rate; blue dots) per condition (*n* = 20). **c** Each trial consisted of a short periodic stream (<10 s) of syllables. Participants performed a Yes/No task and indicated the presence or absence of a perceptual deviant (higher-pitched syllable) at the end of each trial. **d** Average performance (Accuracy; blue dots) per condition (*n* = 15). The same conventions as in Fig. 1 b-d.

In Experiment 6, the comparison of performance (Hit minus FA rate) across different tempi revealed significant differences (repeated-measures ANOVA: F(6,102) = 6.83, p < 0.001; Fig. 5b), with an inverse U-shaped pattern (second-order log polynomial function; adjusted r2 = 0.85) peaking at 1.92 Hz.

Conversely, in Experiment 7, the comparison of accuracy rates across tempi did not reveal significant performance fluctuations (repeated-measures ANOVA: F(6, 84) = 2.75, p = 0.041; Fig. 5d). In this case, there is no discernible optimal frequency.

This indicates that the type of stimulus (tones, syllables) does not have an influence on which attentional mode is operational, only the stream length does.

### Absence of rhythmic mode of operation in syllable categorization tasks

In Experiments 8 and 9, we again employed syllables to test a higher level of processing, this time phonological, by asking participants to detect a specific syllable /ba/ within a stream of diverse syllables (Fig. 6a and Fig 6c; see Methods). The average difficulty level (voice onset time duration) was 16.05 ms (Mdn = 10.5, SD = 13.81) for Experiment 8 (Supplementary Fig. 6a) and 23.5 ms (Mdn = 23, SD = 14.36) for Experiment 9 (Supplementary Fig. 6c).

**Fig. 6:**
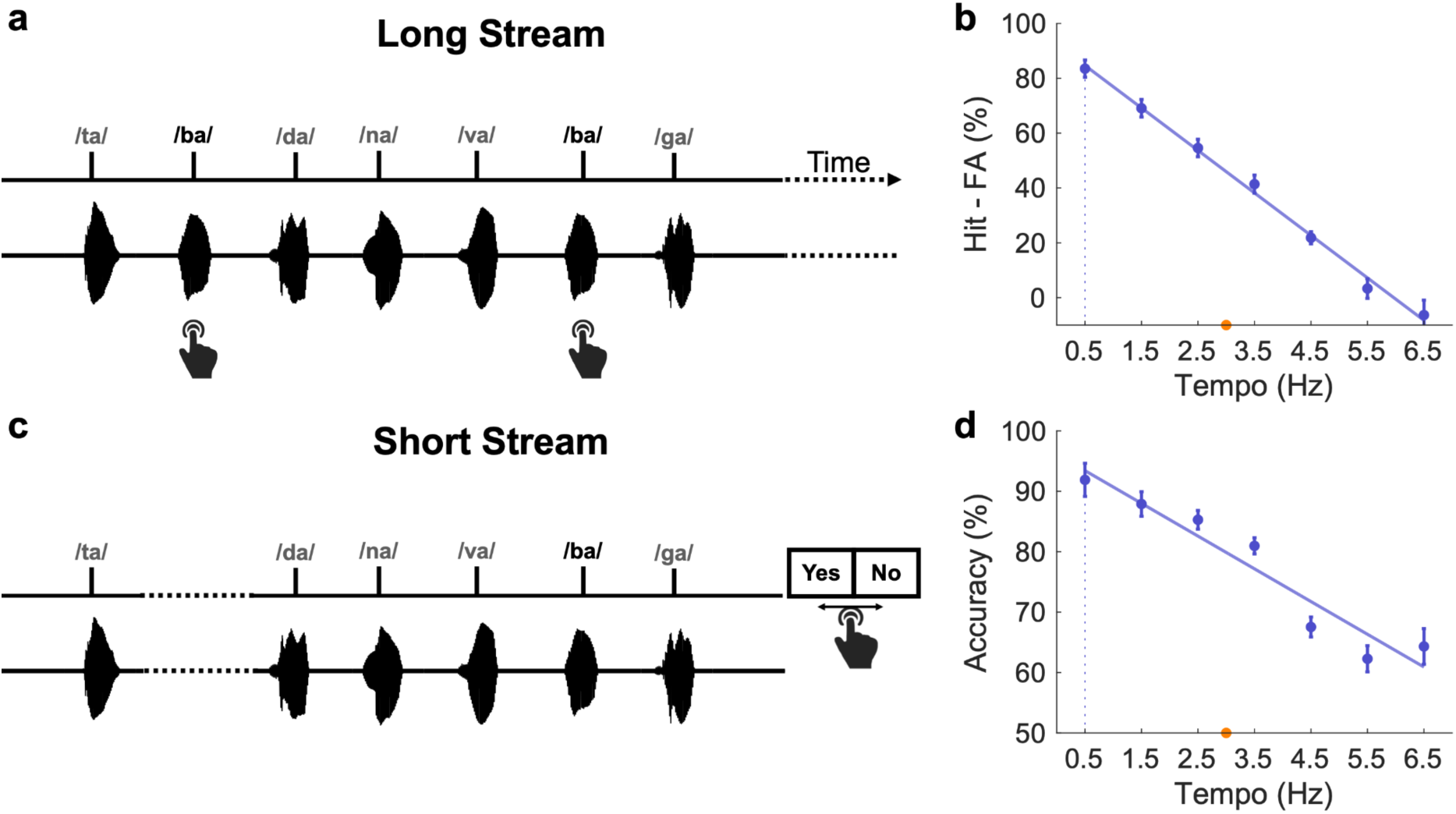
Experiments 8-9. Categorization of syllable /ba/. **a** Each trial consisted of a long periodic stream (∼1 min) of syllables. Participants performed an ongoing categorization task of the target syllable /ba/. **b** Average performance (Hit minus FA rate; blue dots) per condition (*n* = 20). **c** Each trial consisted of a short periodic stream (<10 s) of syllables. Participants performed a Yes/No task and indicated the presence or absence of syllable /ba/ at the end of each trial. **d** Average performance (Accuracy; blue dots) per condition (*n* = 14). The same conventions as in Fig. 1 b-d.

In experiment 8, comparing the performance (Hit minus FA rate) across tempi revealed significant differences (repeated-measures ANOVA: F(6,114) = 76.65, p < 0.001; Fig. 6b). The performance linearly decreased with an increase in frequency and could be approximated by a first-order linear function (adjusted r2 = 0.99) peaking at the slowest presented tempi, 0.5 Hz.

In experiment 9, we observed similar results. The performance (accuracy) showed significant fluctuations (repeated-measures ANOVA: F(6,78) = 27.82), p < 0.001; Fig. 6d) and could be approximated by a first-order linear function (adjusted r2 = 0.91) peaking at 0.5 Hz.

Hence, for categorization tasks, on the one hand, the response profile is similar between the short and long-stream paradigms, and on the other hand, no rhythmic mode of attention is observed (neither around 1.5 nor 4.5 Hz). This indicates that for categorization tasks, the longer the time between two stimuli the better the performance, which is suggestive of a profile of evidence accumulation^37–39^.

### Control analysis

Finally, we performed a control analysis to first confirm that the task type (ongoing detection or Yes/No) has no impact on our results, and second, to better evaluate the dynamics of the emergence of the rhythmic mode of operation. We reasoned that in tasks involving long streams, the rhythmic mode should be absent in the early period and emerge after some time. To analyze the data in a time-resolved manner (corresponding to the moment of occurrence of the 8 different deviant targets within the stream) while keeping a decent statistical power, we regrouped the data collected from all the ongoing detection tasks with long streams (experiments 1, 4, and 6; n = 61). The profile of performance across tempi (flat or inverse U-shaped) was found to be dependent on the moment at which the deviant appeared in the auditory stream (Fig. 7). Performance (Hit rate) was not significantly different across tempi for the first deviant (repeated-measures ANOVA: F(6,360) = 1.53, p = 0.19; Fig. 7a), which appeared, on average, 4.91 s after stimulus onset (Fig. 7b). On the contrary, performance was significantly different across tempi for all the other, subsequent deviants (all F(6,360) > 8.97; all p < 0.001), starting from the second deviant, which appeared on average at 10.25 s. For these latter, the performance profile across tempi exhibited an inverse U-shaped pattern (second-order log polynomial function; all adjusted r2 > 0.87) peaking between 2.06 to 2.43 Hz.

**Fig. 7:**
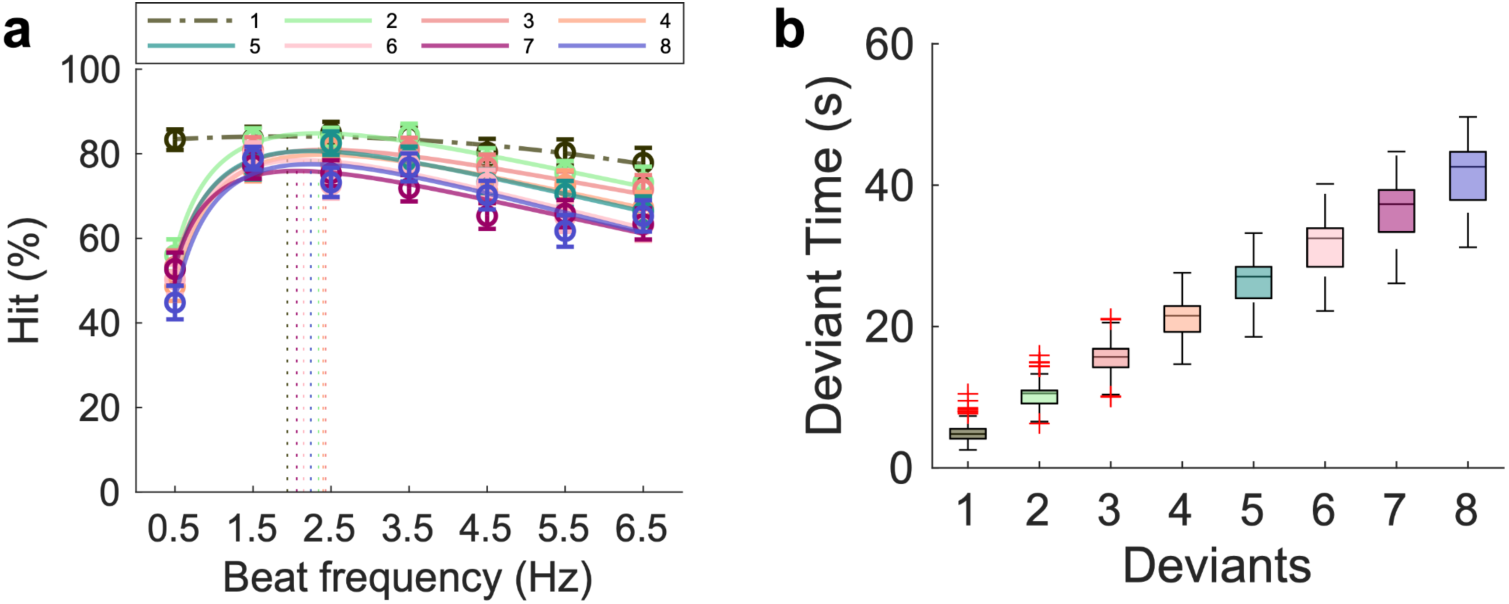
Control analyses. **a** Average performance across all the ongoing detection tasks (experiments 1, 4, 6) per condition (*n* = 61), decomposed as a function of the index of the deviant in the stream. If performance significantly differed between conditions (repeated-measures ANOVA), data were approximated with a polynomial function (plain line) and an optimal frequency (leading to a maximal performance) could be estimated (vertical dashed line). Else, data were approximated with a polynomial function (dashed line). Error bars indicate ±SEM. **b** Each box plot represents the distribution of moment of occurrence (in seconds) of each of the eight deviants, across trials, conditions, and experiments, with the central line marking the median, the box edges the quartiles, and the whiskers the non-outlier edge.

## Discussion

In this comprehensive study, through a series of 9 behavioral experiments, we explored the existence of two modes of attention and examined the factors that determine when they are deployed. We explored a range of variables, including the length of auditory streams, perceptual features (such as duration and pitch), types of stimuli (such as pure tones or syllables), the presence or absence of distractors in a stream, and the specific task at hand, namely whether it was perceptual or categorical. Our experiments provide behavioral evidence for two distinct attentional modes^4^. Initially, the continuous mode is predominant upon the introduction of a stimulus, but it transitions to a rhythmic mode after approximately 7 s (Fig. 7) and is characterized by an optimal sampling rate of approximately 1-2.5 Hz^20^ (Fig. 1b, 3b, 4b, 5b). Fascinatingly, the nature of the task determined these modes of attention, as they emerge solely in perceptual tasks. Conversely, in categorization tasks, we observe a robust linear decrease in performance as the tempo increases (Fig. 6b, 6d). A model of coupled oscillators captures these perceptual effects and indicates that the transition to a rhythmic mode of operation is related to a progressive decrease in the noise level in the system (Fig. 2 and Supplementary Fig. 2).

A popular hypothesis suggests that attention operates in rhythmic or continuous modes depending on the dynamics of task demands^4,28^. This suggests an adaptive attentional framework capable of optimizing sensory processing and behavioral performance by dynamically adjusting its operational mode in response to environmental demands. While previous research has investigated the neural bases and behavioral consequences of a rhythmic mode of perception^40–43,19,20^, no study to our knowledge has provided direct behavioral evidence of these two modes, and of how they transition between one another. In our study, the pivotal finding is the crucial role of auditory stream length in determining the shift between different attentional modes. Specifically, long auditory streams, those extending beyond 5 - 7 s, consistently triggered a transition from a continuous to a rhythmic mode of attention. This finding is consistent with prior research, which demonstrated that brain activity synchronizes with the stimulus’s temporal structure, particularly for longer streams^15–17,24,44–47^. It also explains why other studies —using short streams— did not observe perceptual benefits with rhythmic streams^26,27^, reconciling apparent contradictory findings under a common framework.

Our findings indicate that upon introducing a stimulus stream, there is an inclination towards engaging in a continuous mode of attention, which can be maintained for approximately 7 s, irrespective of the tempo, the perceptual feature to detect, or the nature of the stimulus. Hence, while it was proposed that the continuous mode operates for temporally unpredictable sensory streams^4^, we show that it also operates for a few seconds after the onset of highly predictable temporal structures. This continuous mode of operation is characterized by a high accuracy across all tempi. This mode is suggested to facilitate rigorous responses to all presented stimuli and is presumably more energy-intensive^4,48,49^, thereby rendering it sustainable for only a limited timeframe. Consequently, this continuous vigilance mode is circumvented in favor of a more optimized, rhythmic mode of processing, if the task and the stimulus dynamics allow it. This strategic shift would be a solution for the constraints of cognitive resources over extended periods, highlighting the necessity for modulation of attention to maintain performance accuracy. For this rhythmic mode to operate, the stimulus dynamics must be periodic, temporally predictable, and occur at the right tempo. Indeed, our results confirm that the rhythmic mode is optimal at approximately 1.5 Hz in the auditory domain^20^ (1-2.5 Hz range). According to the active sensing framework, the dynamics of the motor cortex influence auditory perception^50^, thereby forming an action-perception closed-loop system^51,52^. This indicates that the dynamics of the motor system are instrumental in shaping the temporal organization of sensory systems, thus affecting the processing of sensory inputs^52^. Thus, it is plausible that in an experimental context involving active tracking—rather than passive listening—the transition from a continuous to a rhythmic mode may occur more quickly. The observed frequency range of 1-2.5 Hz not only aligns with the rate of natural or precise human rhythmic movements such as finger tapping, but also matches with the rate (0.8-2.5 Hz) at which our strongest sense of music beats occurs^21,22,53–56^. This frequency range may be linked to the contribution of motor-related cortical and subcortical structures^57–61^. The influence of the motor system on auditory perception has been shown for both music^35,36,62^ and speech^63,64^. Specifically, in the context of music, studies involving beat-based perception tasks^20,55,56^ indicate that such structures are pivotal for making temporal predictions in auditory attention contexts^36,52^ and operate within certain rate limits^20,52^, particularly evident in the beta (20 Hz) and delta (1.5 Hz) neural dynamics^20,36,52^. Alternatively, such preference may reflect a physiological constraint imposed by the time constant of neural dynamics of auditory subcortical structures^65,66^ or the auditory dorsal pathway^67^.

Similarly, in speech, there is a convergence of evidence highlighting the critical role of motor cortical dynamics, particularly within the delta and theta frequency range. Spontaneous finger tapping to the perceived prosodic rhythm of speech typically occurs within the delta range^68,69^. The tracking of acoustic dynamics by the left auditory cortex is primarily modulated by motor areas through delta (and to some extent, theta) neural dynamics^63,70^. Furthermore, delta-tracking of the phrasal acoustic rate and delta-beta coupling in motor areas predicts speech comprehension^64^. Additional studies identify an optimal frequency for speech perception and intelligibility around 4 to 6 Hz, within the theta range^71–74^. Additionally, the strongest audio-motor coupling has been shown for the presentation rate of 4.5 Hz^32^, aligning closely with the average syllabic rate across most languages^5,31,32^. Despite these insights, whether speech and music perception exhibit distinct optimal rates has not been extensively studied using comparable methodologies. In this study, using similar paradigms, we show an optimal performance accuracy at 1.5 Hz in perceptual tasks involving streams of pure tones and syllables. It is important to note that in all the previous tasks involving syllables, no behavioral effect of audio-motor coupling was observed at a syllabic rate of 4-5 Hz^31–33,75^. Furthermore, the lack of difference in response to various stimuli directly contrasts with findings suggesting distinct rate-specific temporal processing mechanisms for detection tasks involving speech and music stimuli^75^. Our findings suggest instead a generalized auditory periodic temporal attention processing framework for perceptual tasks.

To further elucidate the dynamic transition between a continuous and rhythmic mode of operation, we employed a simple model of coupled oscillators (Fig. 2a). Adjusting the noise parameter in the model was critical to fit the observed experimental outcomes. It is possible that noise plays an important role in facilitating the emergence of synchronization^76^; however, over time, as synchronization becomes more robust, the system may reduce the impact of noise^77^. Through noise level adjustment, this model provides a potential mechanistic explanation for the observed differences in outcomes between short and long auditory streams. However, this model is simplistic, focusing solely on the synchronization of an attention oscillator to a periodic stimulus. Further developments are necessary for the model to accurately capture the complexities associated with the two modes of attention and different task types. The precise duration necessary to initiate this shift remains to be determined through further research. Still, control analyses suggest that the rhythmic mode needs approximately 5 - 7 s to emerge (Fig. 7). It is noteworthy that empirical studies employing a musical prime to establish the rhythmic attentional mode and enhance subsequent language processing typically utilize primes with a long duration, of approximately 30 s^78–82^. Similarly, when listening to long auditory streams, lapses of attention tend to occur at a rate of 0.06 Hz, corresponding to a period of approximately 15 s^41,83^. Stimulus-brain phase alignment also dynamically evolves during at least the first 8 s of stimulation, which suggests that the brain progressively attunes to external rhythmic stimulation by adapting its internal representation of incoming environmental stimuli^84^. Finally, this rhythmic mode of operation does not last forever, and while highly rhythmic musical stimulation facilitates perception for at least 3 min, longer exposure (9 min) ends up reducing the sensitivity of the auditory system^85^.

An additional finding in our study is the absence of evidence for either rhythmic or continuous mode in tasks involving categorizing the syllabic category /ba/, across both long continuous and short streams of syllables. A linear decrease in accuracy with increasing presentation frequency was observed, indicating that slower presentation rates allow more time for processing and accurate syllable categorization. This set of results aligns with the traditional evidence accumulation model^38^, indicating that performance is compromised when there is insufficient time to reach the decision threshold.

In conclusion, our study provides behavioral evidence for the existence of two modes of attention for perceptual tasks and elucidates the factors that influence their transition. By demonstrating a general mechanism across various auditory tasks and highlighting the critical role of stream length in attentional mode switching, our research advances the understanding of the modes of attention. The absence of mode-specific effects in syllable categorization tasks adds another layer to our understanding, highlighting the strong impact of the processing stage on performance accuracy. Our findings lay the groundwork for future research into the neural mechanisms behind these attentional modes, their consistency across various sensory modalities, and the interaction between auditory and motor systems.

## Methods

### Participants

21, 21, 20, 20, 20, 20, 15, 20, and 14 participants (age range: 18 - 44 years; 71.4% of females) were respectively recruited for experiments 1 to 9. All participants had normal hearing and no neurological or psychiatric disorder history and provided informed consent according to the local ethics committee guidelines from Aix-Marseille University and Aarhus University.

### Overview of experiments

This study comprises nine behavioral experiments examining various factors like stream length, stimuli type, attended features, tasks, and tempi as summarized in Supplementary Table 1.

### Stimuli

Auditory stimuli were presented at a sampling rate of 44.1 kHz using the Psychophysics Toolbox^86^ within Matlab (MathWorks) for all experiments.

For experiments 1-5, pure tones were created. Each standard tone was 80 ms long with a dampening length of 10 ms and a 40 dB attenuation.

For experiments 6-9, syllables were recorded from a single speaker in a soundproof room through a Realtek sound card and edited using Audacity (The Audacity team). Each standard syllable was 140 ms in length and all syllables had a 15 ms linear offset ramp. For experiments 6 and 7 (higher-pitch target syllables), five standard syllables (/da/, /ga/, /pa/, /ta/, and /va/) had a fundamental frequency of 250.5 Hz. Pitch variants of syllables /pa/, /ta/, and /va/ were generated for the staircase. Their frequencies ranged from 250.5 Hz - 275.6 Hz. For experiments 8 and 9 (categorical target /ba/), three versions of each syllable differing in pitches were recorded for each of the standard syllables (/da/, /ga/, /na/, /ta/, /va/, and /pa/). However, the third version was removed for /da/, /ga/, and /na/ due to prominent artifacts.

For Experiments 8 and 9, /ba/ stimuli were generated and edited for use in the staircase procedure by reducing the voice onset time (VOT) from 60 ms to 0 ms. In Experiment 8, twenty /ba/ stimuli were created with VOTs varying in 3 ms increments using an onset ramp. In Experiment 9, sixteen /ba/ stimuli were generated with VOTs varying on a logarithmic scale.

### Experimental design

All experiments presented stimuli binaurally through headphones (Sennheiser HD 250 linear) with a 44.1 kHz sampling rate. Volume was kept at a constant comfortable level throughout all experiments. Before each experiment, the difficulty level was determined (to avoid the ceiling or floor effects) using an adaptive staircase procedure. This also served to familiarize participants with the stimuli and task. For the staircase procedure, participants were presented with auditory streams at a rate of 3 Hz, except, in experiment 3, where the streams were presented at 1.5 Hz. During the main experiment, participants were presented with sequences at seven tempi (0.5, 1.5, 2.5, 3.5, 4.5, 5.5, and 6.5 Hz) for experiments 1, 2 and 4-9 and five tempi (0.7, 1, 1.5, 2.3 and 3.5 Hz) for experiment 3.

For experiments 1, 4, and 6, participants performed an ongoing detection task of detecting perceptual deviants embedded in long auditory streams. These deviant targets varied per experiment: a higher-pitched tone for experiment 1, a tone with a longer duration for experiment 4, and a higher-pitched syllable for experiment 6. Each auditory stream embedded 8 (experiment 1 and 4) or 9 (experiment 6) deviants. There were 6 blocks in total, with each presentation rate occurring once per block, in random order. As a result, there were 48 or 54 deviants for each presentation rate. Participants were told to not move during trials and to press the down-arrow key as fast as possible when they heard the deviant.

For experiments 2, 5, and 7, participants performed a Yes/No task. Each trial consisted of a short stream of auditory stimuli and the participants had to evaluate if a single perceptual deviant was present in the stream or not. Each experiment involved a different deviant target: a higher-pitched tone for experiment 2, a tone of longer duration for experiment 5, and a higher-pitched syllable for experiment 7. At the end of each trial, participants had to press the right-arrow key if they had heard the deviant in the stream, or else the left-arrow key. There were 21 blocks in total, with 14 trials of the same presentation rate per block, which resulted in 42 trials for each presentation rate.

For experiment 3, participants performed a pitch evidence-accumulation task. Each trial consisted of 15 pure tones: Three reference tones preceded an alternation of six targets and six distractor tones of variable frequencies. Targets occurred in phase with the preceding references, whereas distractors occurred in antiphase so that participants could use the beat provided by the references to distinguish targets from distractors, which were otherwise perceptually indistinguishable. Participants performed a pitch evidence-accumulation task at the end of each trial, by pressing the up (/down) arrow key with their right hand if they considered that the mean frequency of targets was higher (/lower) than the reference frequency (f0). The mean frequency of distractors was always equal to f0, and hence noninformative. The absolute difference between the frequency of targets and f0 was titrated for each participant to reach 75% of categorization performance (psychophysical staircase procedure). Only five tempi were investigated in this experiment (see above). After a short training and the staircase procedure, participants performed 10 blocks in total, with 24 trials of the same presentation rate per block, which resulted in 48 trials for each presentation rate.

In Experiment 8, participants engaged in an ongoing categorization task, similar to those in experiments 1, 4, and 6, where they were asked to detect the target syllable /ba/. Each long auditory stream contained 9 embedded instances of the target syllable. In Experiment 9, participants completed a Yes/No task, similar to those in experiments 2, 5, and 7, where they evaluated whether the syllable /ba/ was present in a short auditory stream.

For all experiments, participants were provided feedback indicating the number of correct responses and the number of false alarms and were encouraged to take breaks between trials. All experiments lasted approximately one hour.

### Data processing and statistical analyses

For long-stream ongoing detection/categorization tasks (experiments 1, 4, 6, and 8), button presses that occurred 0.25 to 1.2 s^19^ following the target were considered hits, and presses outside this window were counted as false alarms. Hit rate (%) and false alarms (%) were analyzed as dependent measures as the variable number of standards per presentation rate precluded the analysis of d’.

For short streams, Yes/No tasks (experiments 2, 5, 7, and 9) and experiment 3, reaction time (RT) outliers were detected using the median plus or minus 2.5 times the Median Absolute Deviation (MAD) method^87^. Trials with RT outliers were removed during pre-processing. Responses with correct identification of a target were counted as hits, and other responses were considered false alarms. Participants’ responses were z-scored to calculate the d’ (zHit - zFA).

All analyses were performed in MATLAB (MathWorks) at the single-subject level and followed by standard parametric two-sided tests (repeated-measures ANOVAs) at the group level. The threshold for statistical significance was defined at p < 0.01. When significant differences across tempi were observed, the optimal tempo of performance was determined by testing different models on subject-averaged data: Variations of performance across tempi were approximated by either first, second, or third-order polynomial functions and on either a linear or log-scaled x-axis (tempi) resulting in six models. The best-fitting model was determined using the maximal adjusted R-squared value.

### Model of coupled oscillators

We implemented a model of two-coupled phase oscillators^88^ with time delays^89^, time-varying noise, and time-varying coupling strength to approximate the selective coupling between a periodic external beat (stimulus; S) and auditory periodic temporal attention (A) (Fig. 2a). The model was implemented with a set of differential equations, as:

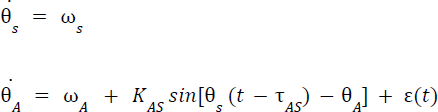

Where ω_i_, *θ_i_* and ε*_i_* are the natural frequency, phase, and noise of oscillator i, and for each pair of oscillators i and j, *K_ij_* and *τ_ij_* represent the coupling strength and time-delay from oscillator i to j. By default, the noise was set to additive and Gaussian with an intensity value of 10, the coupling strength was set to 13, and the time delay to 0.1s^20^.

Additionally, the noise, coupling strength, and time-delay parameters were also varied with time, in three different versions of the model. The noise was varied using a reverse sigmoid function:

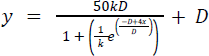

Where k is a constant, set to 1.5.

The coupling strength was varied linearly as: *y* = 0. 265*x*

The time delay was varied linearly as: *y* = 0. 4 − 0. 006*x*

The sampling rate of the simulation was 25 ms. We set internal time delays to 0 ms (i.e., < 25 ms). The level of coherence between the external beat (S) and the auditory attention oscillator (A) was computed with the phase-locking value (PLV). It estimates the capacity of

A to synchronize S and is hence used as an approximation of behavioral performance. PLV is defined as:

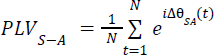

Where the phase angle Δθ between oscillators S and A at time t is averaged across time points from t = 1 to N.

## Code availability

All code used for all analyses and plots will be publicly available on GitHub (https://github.com/orgs/ins-amu/teams/dcap) following the manuscript’s publication.

## Acknowledgments

We thank Arnaud Zalta for his contribution to Experiment 3, Céline Hidalgo for syllable recordings, and all the members of D-CAP-INS for helpful discussions throughout this project. This study was supported by ANR-20-CE28-0007 (to B.M), ERC-CoG-101043344 (to B.M), Fondation Pour l’Audition (FPA RD-2022-09; to B.M.), and grants from France 2030 (ANR-16-CONV-0002) and the Excellence Initiative of Aix-Marseille University (A*MIDEX AMX-19-IET-004).

## Author contributions

All authors conceived the experiment(s), B.M. supervised the work, A.G. and T.M. collected the data, A.G. and B.M. wrote the analysis code and analyzed the data, A.G. wrote the first draft of the manuscript, and all authors edited the manuscript.

## Supplementary Figures

**Supplementary Fig. 1:**
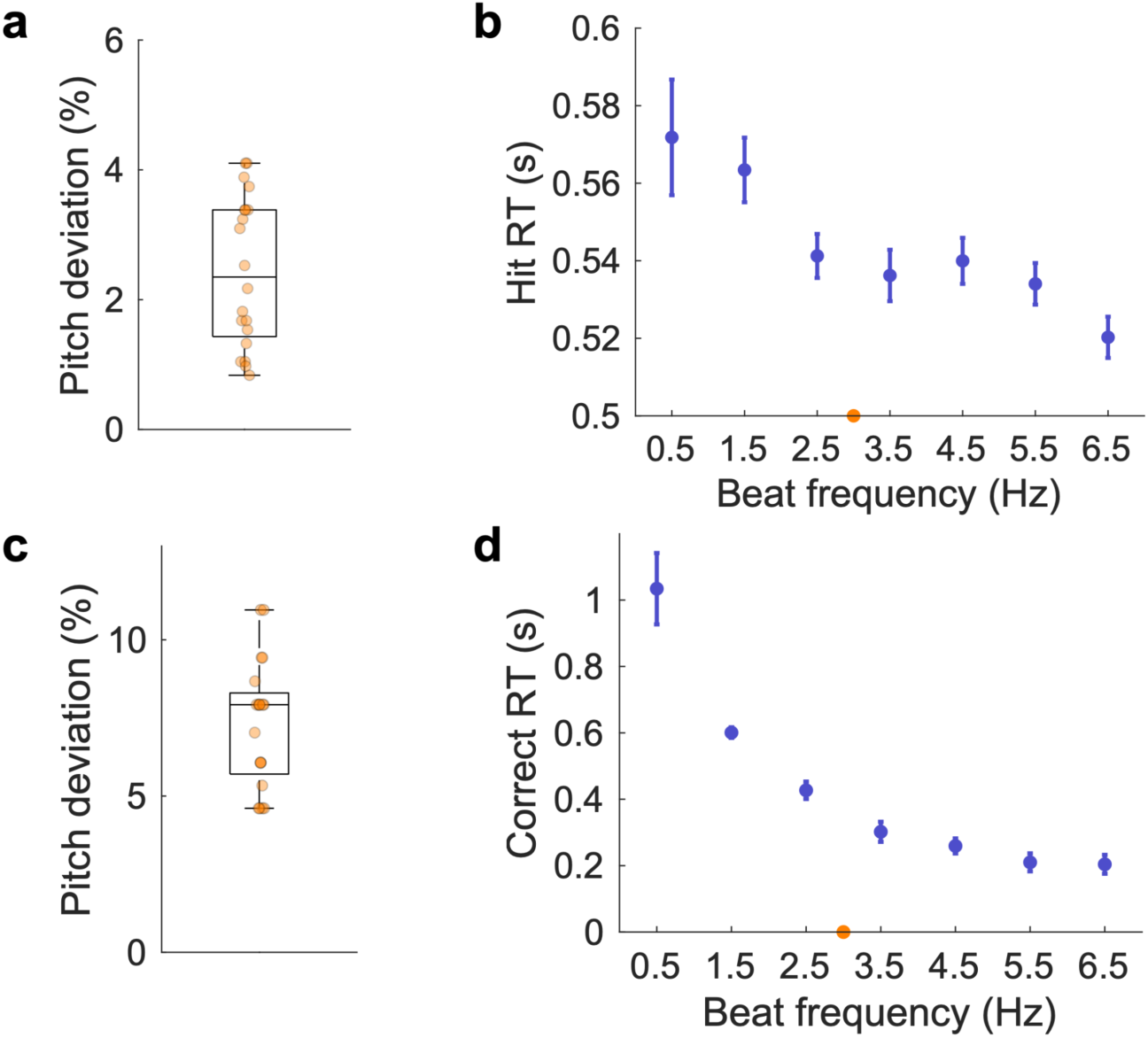
Experiments 1-2. Detection of higher-pitched tones. **a, c** An adaptive staircase procedure defined the individual difficulty level to reach threshold performance for a 3 Hz tempo for Experiment 1 (**a**) and Experiment 2 (**c**). The difficulty was modulated by adjusting the pitch of the deviants (in percent compared to standard tones (f_0_=660 Hz), estimated in base-2 logarithmic units). The boxplot represents the median and 1.5 times the interquartile range. **b, d** Average hit (**b**) or correct (**d**) reaction times (RT) per condition for Experiment 1 (**b**, *n* = 20) and Experiment 2 (**d**, *n* = 21). The orange dot indicates the tempo (3 Hz) that served to assess the individual difficulty level (**a, c**). Error bars indicate ±SEM.

**Supplementary Fig. 2:**
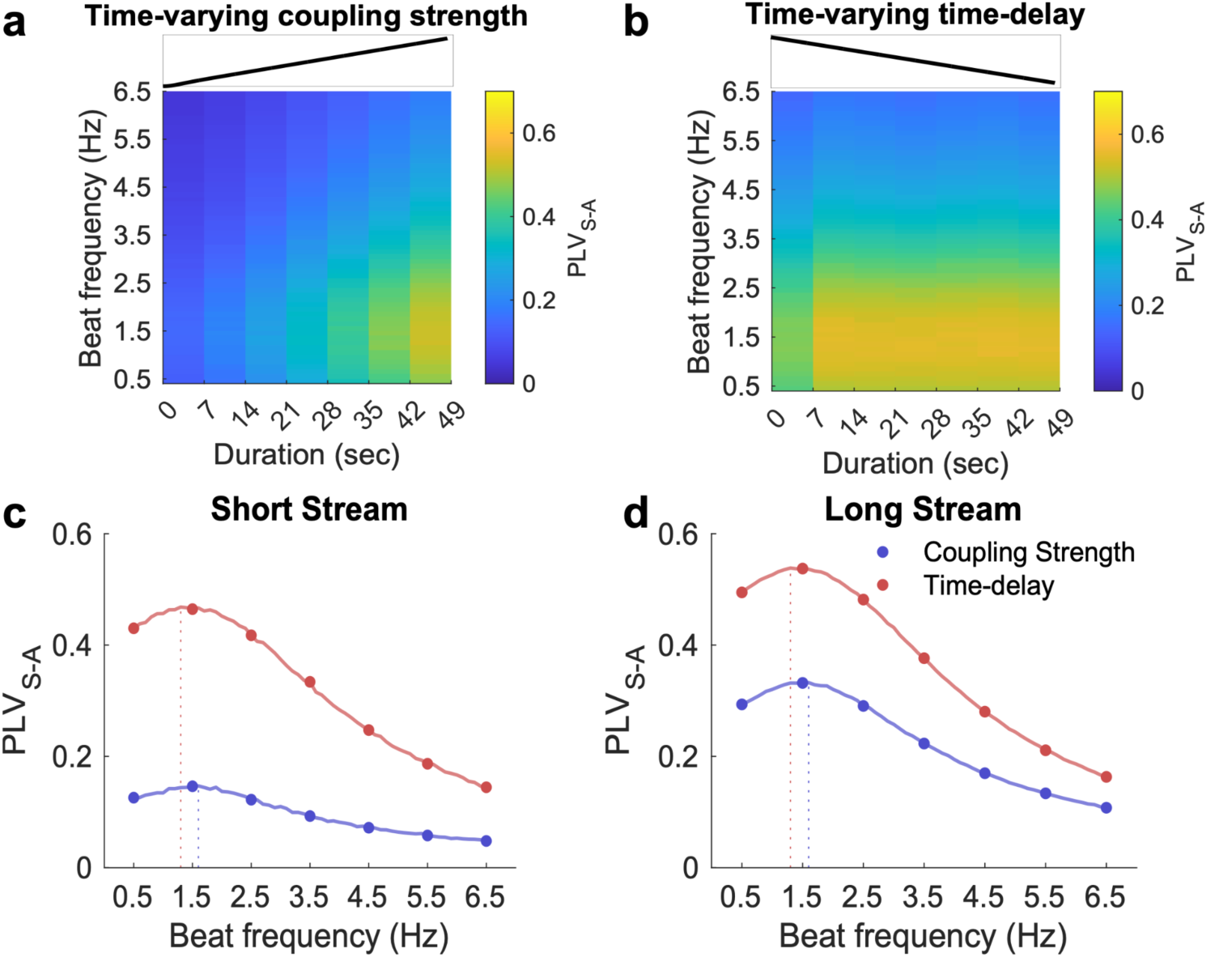
A model of coupled oscillators to generalize the results. **a, b** Heatmap depicting the evolution of PLV for each condition (tempi), from 0 to 49 s, in steps of 7 s. The tempi varied from 0.5 Hz to 6.5 Hz (in 0.01 Hz steps) and data of n = 20 participants were simulated. The **(a)** coupling strength K or **(b)** the time-delay τ varied over time (see insets) while the other parameters, noise N and **(a)** K or **(b)** τ were kept constant. **(c, d)** Time-averaged PLV corresponding to **(c)** short (0-7 s) or **(d)** long (0-49 s) trials for time-varying coupling strength (blue) and time delay (red) variations over time. The same conventions as in Fig. 2 c-d.

**Supplementary Fig. 3:**
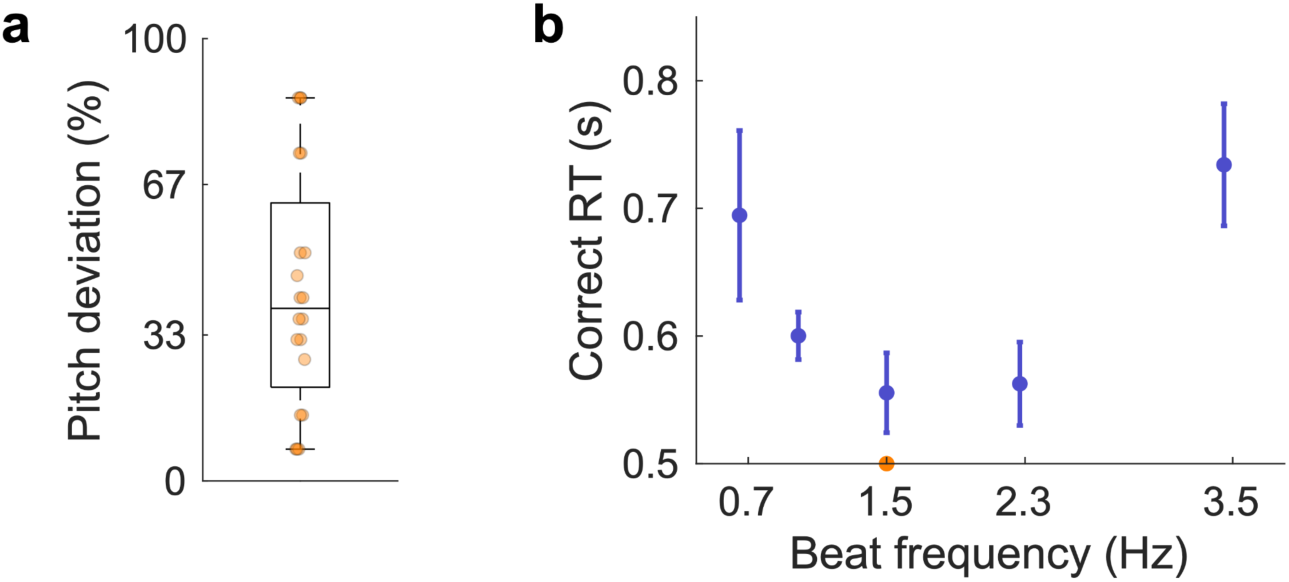
Experiment 3. Pitch evidence-accumulation. **a** Individual difficulty level to reach threshold performance for a 1.5 Hz tempo. The difficulty was modulated by adjusting the absolute difference between the frequency of targets and the reference frequency (f0). **b** Average correct RT per condition (*n* = 20). The same conventions as in Fig. S1.

**Supplementary Fig. 4:**
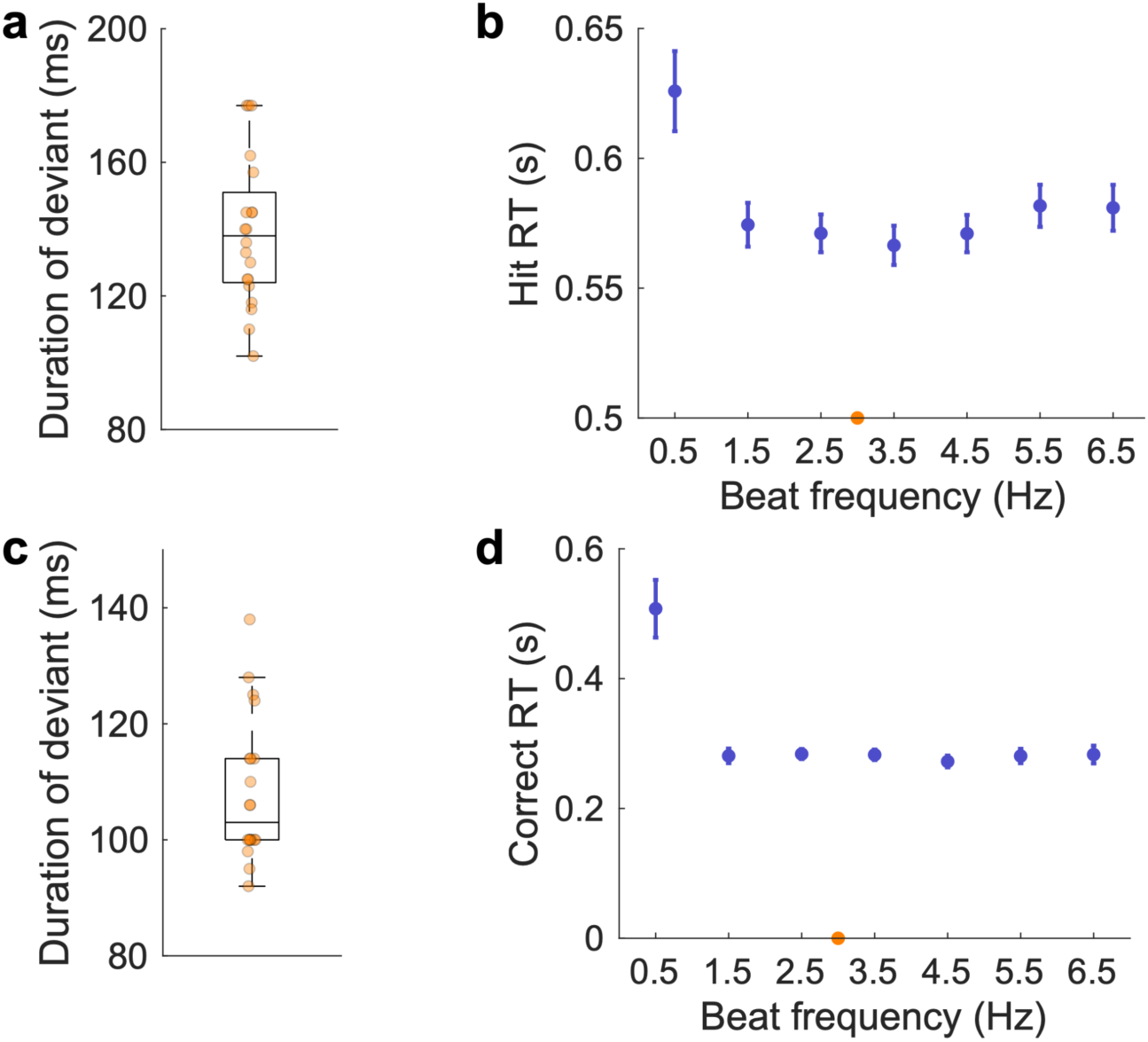
Experiments 4-5. Detection of longer tones. **a, c** Individual difficulty level to reach threshold performance for a 3 Hz tempo for Experiment 4 (**a**) and Experiment 5 (**c**). The difficulty was modulated by adjusting the duration of the deviants (standard tones had a length of 80 ms). **b, d** Average hit (**b**) and correct (**d**) RT per condition for Experiment 4 (**b**, *n* = 20) and Experiment 5 (**d**, *n* = 20). The same conventions as in Fig. S1.

**Supplementary Fig. 5:**
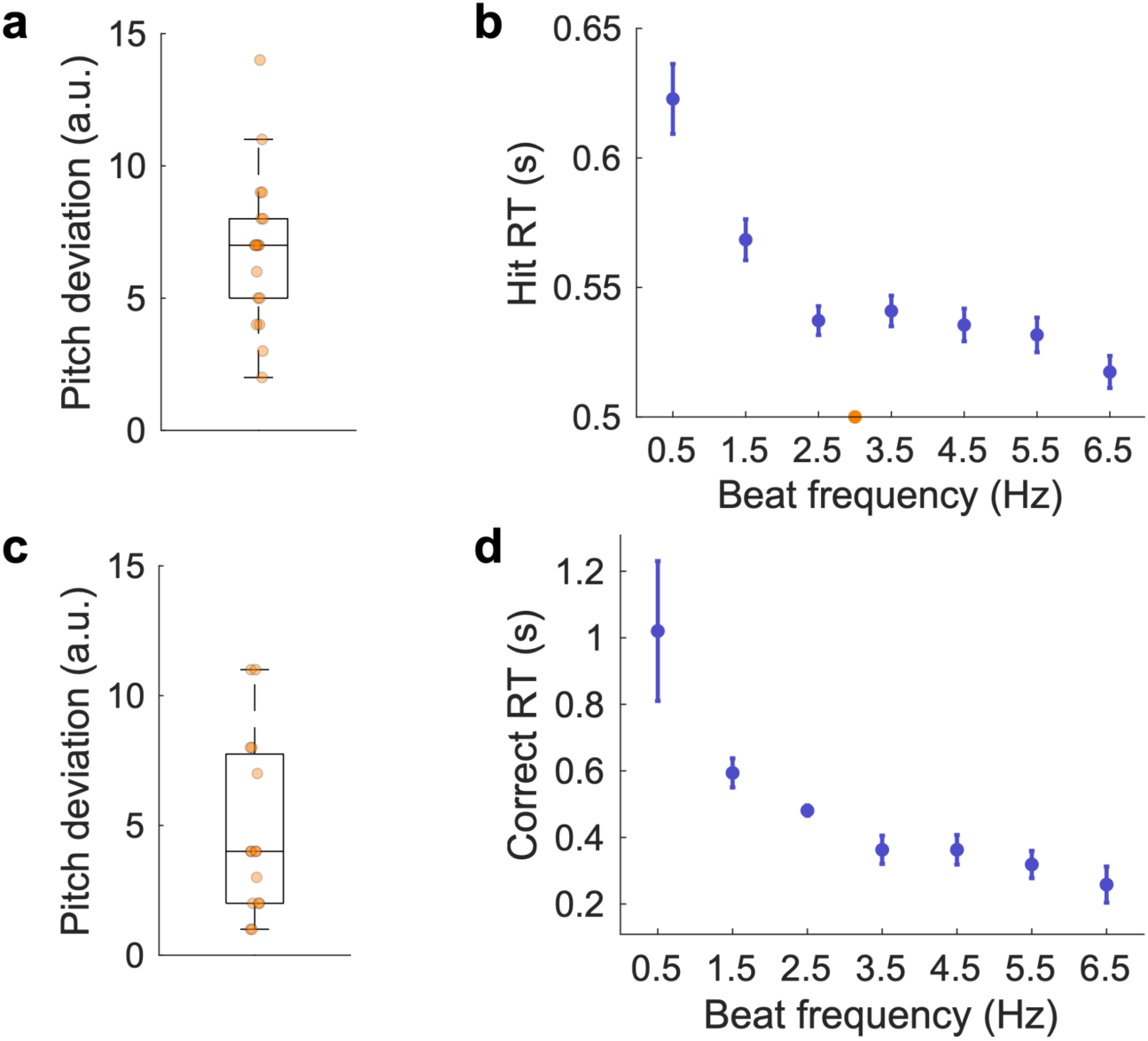
Experiments 6-7. Detection of higher-pitched syllables. **a, c** Individual difficulty level to reach threshold performance for a 3 Hz tempo for Experiment 6 (**a**) and Experiment 7 (**c**). The difficulty was modulated by adjusting the pitch of the deviant syllables. **b, d** Average hit (**b**) and correct (**d**) RT per condition for Experiment 6 (**b**, *n* = 20) and Experiment 7(**d**, *n* = 15). The same conventions as in Fig. S1.

**Supplementary Fig. 6:**
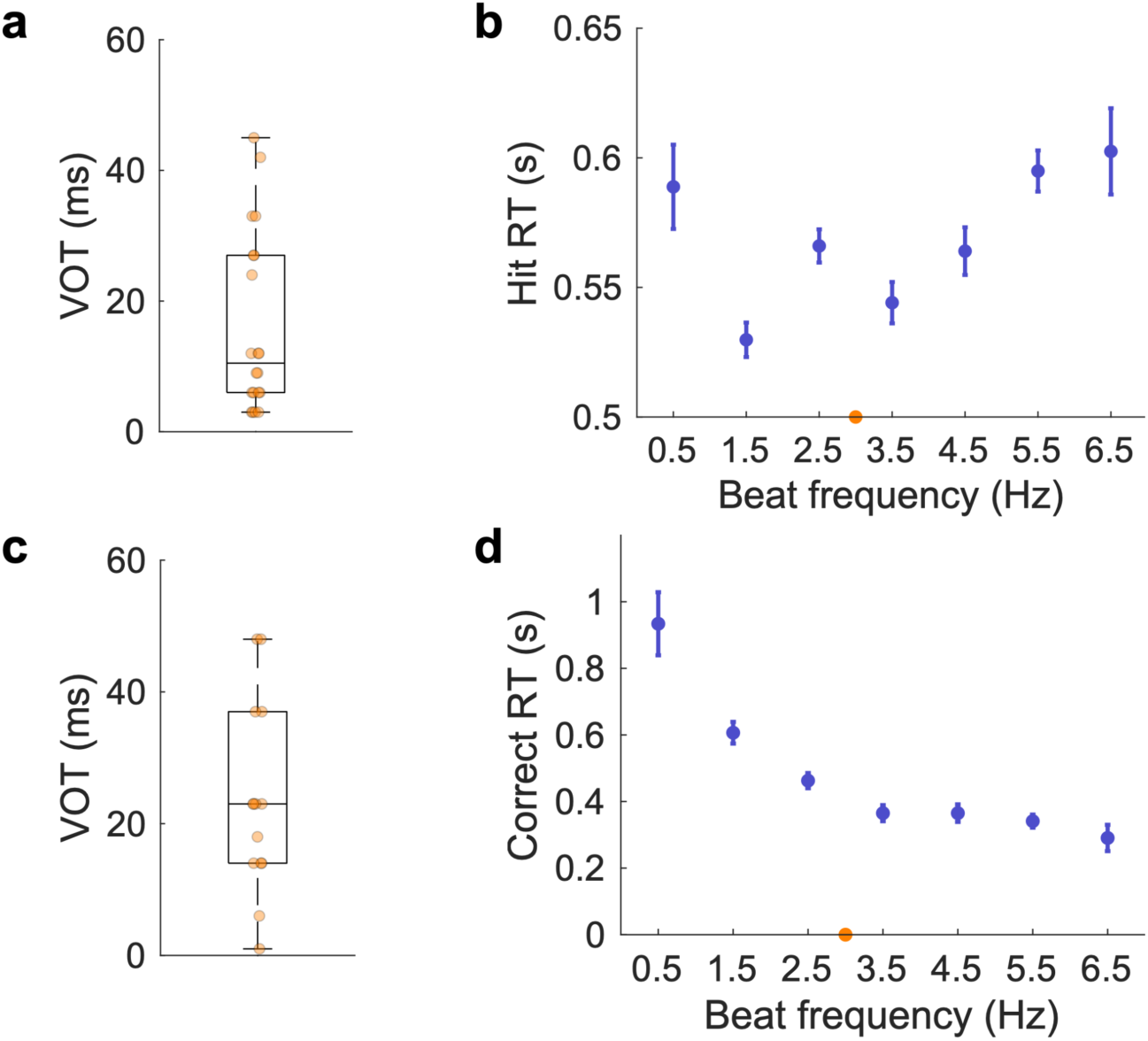
Experiments 8-9. Identification of syllable /ba/. **a, c** Individual difficulty level to reach threshold performance for a 3 Hz tempo for Experiment 6 (**a**) and Experiment 7 (**c**). The difficulty was modulated by adjusting the VOT of the syllable /ba/. **b, d** Average hit (**b**) and correct (**d**) RT per condition for Experiment 8 (**b**, *n* = 20) and Experiment 9 (**d**, *n* = 14). The same conventions as in Fig. S1.

## Supplementary Tables

**Supplementary Table 1.** Experiments Summary. Table summarizing the length of stimulus, type of stimulus, type of task, and the feature tested for all experiments conducted in the study.

| Experiment | Stream Length | Stimuli Type | Task Type | Attended Feature |
| --- | --- | --- | --- | --- |
| 1 | Long | Pure tones | Detection | Higher-pitched tones |
| 2 | Short | Pure tones | Yes/No | Higher-pitched tones |
| 3 | Short | Pure tones | Evidence Accumulation | Pitch detection |
| 4 | Long | Pure Tones | Detection | Longer duration tone |
| 5 | Short | Pure Tones | Yes/No | Longer duration tone |
| 6 | Long | Syllables | Detection | Higher-pitched syllable |
| 7 | Short | Syllables | Yes/No | Higher-pitched syllable |
| 8 | Long | Syllables | Identification | Syllable /ba/ |
| 9 | Short | Syllables | Yes/No | Syllable /ba/ |

